# Memory consolidation during rest forms shortcuts in a cognitive map

**DOI:** 10.1101/2024.01.22.576595

**Authors:** Cal M. Shearer, Annalise B. Rawson, Helen C. Barron, Jill X. O’Reilly

**Affiliations:** Department of Experimental Psychology, University of Oxford, Oxford, United Kingdom; Medical Research Council Brain Network Dynamics Unit, Nuffield Department for Clinical Neurosciences, University of Oxford, Oxford, United Kingdom; Wellcome Centre of Integrative Neuroimaging, University of Oxford, FMRIB, John Radcliffe Hospital, Oxford, United Kingdom

## Abstract

Rest and sleep not only strengthen existing memories but also reorganise memories to generate new knowledge that extends beyond direct experience. However, it remains unclear both *how* memories are reorganised and the effect of this reorganisation on behaviour. Here, we designed a novel protocol to casually manipulate memory consolidation during rest using awake, contextual targeted memory reactivation (TMR). We found that promoting memory consolidation during rest qualitatively reorganises memories by *forming ‘shortcuts’ between memories* which have not been experienced together. These shortcuts in memory extend beyond direct experience to facilitate our ability to make novel inferences. A series of control tests indicate that inference performance cannot be explained by quantitative strengthening of the experienced component links but are rather explained by qualitative changes in the cognitive map which involve formation of new shortcuts. Interestingly, we show that representing a shortcut may come with limitations, as shortcuts cannot be readily updated in response to rapid changes in the environment. Together, these findings reveal how memories are reorganised during awake rest to construct a cognitive map of our environment, while highlighting the constraints set by a trade-off between efficient and flexible behaviour.

## Introduction

To understand our environment, we construct a cognitive map^1^ forged from associations we have experienced. However, the cognitive map is more than the sum of its parts. As a simple example, if an observer has seen the relationship between A->B and B->C, they can infer the unobserved relationship A->C^2^. It remains unclear when and how these inferential relationships are built to allow us to make predictions that go beyond direct experience. For example, inference could be supported by a chain of learned associations (A->B, B->C), or by formation of a new ‘shortcut’ (A->C). The formation of a new shortcut (A->C) would represent a *qualitative* change in the organisation of memories, which is an essential pre-requisite for the organization of knowledge into a structured cognitive map.

The process of constructing a cognitive map is thought to be facilitated by offline periods of rest/sleep^3–10^. During these periods, memories for previous experiences are reactivated or ‘replayed’^11–13^. Replay involves temporally structured spiking activity that recapitulates previous waking experience^14–16^ to facilitate memory consolidation^11–13,17^ and subsequent decision-making^18^. In addition, growing evidence suggests replay may play a particular role in extending the cognitive map beyond previous waking experience to anticipate upcoming events^19,20^, restructure knowledge^21^, or even “join-the-dots” between spatial trajectories or events that were not directly experienced together^22–24^. However, the way in which memories are reorganised and the consequences of this reorganisation for behaviour remain unknown. Here, we test the hypothesis that offline periods of rest *qualitatively* reorganise memories, rather than *quantitatively* changing the strength of associations that were actually experienced. Further, we predict that this qualitative re-organisation allows for the creation of new inferred links (or shortcuts) that are independent of experienced associations, thus forming a cognitive map that is more than the sum of its parts.

In order to causally manipulate which memories were replayed during offline periods of rest, we developed a novel protocol using awake contextual targeted memory reactivation (TMR). TMR involves playing sensory cues during offline periods of rest or sleep that have previously been linked to a specific memory during prior learning. For directly experienced associations, TMR has been shown to enhance behavioural readouts of memory for cued compared to uncued associations^25–29^. Crucially, TMR has also been shown to bias the content of replay towards cued associations^27^. Given the proposed role of replay in reorganising memory to extend the cognitive map beyond direct experience, and evidence to suggest that TMR can bias the content of replay, we predicted that TMR might provide a tool to investigate how memories are reorganised during periods of rest and the consequences for behaviour. Thus, we sought to test the hypotheses that TMR improves inferential choice and that this improvement is driven by the formation of new inferred links (or ‘shortcuts’) between cues that have not been experienced together^22,30^.

To test our hypothesis, we designed a multi-stage inference task adapted from a protocol previously implemented in humans and mice^22^. In addition, we implemented a novel TMR protocol involving awake, contextual auditory TMR. Consistent with our predictions, we demonstrate that biasing memory reactivation during rest improves the ability to infer novel relationships between sensory cues. Furthermore, using behavioural and physiological (eye-tracking) data, we show that this improvement in inference is driven by the formation of novel shortcuts between indirectly linked cues. As a result of these shortcuts, participants are less reliant on memory of intermediary associations (A->B and B->C) since they can instead use the direct, yet unobserved, shortcut formed during rest (A->C) to inform their decisions. However, although beneficial for inference, we show that building cognitive shortcuts may come with limitations: when the component associations are modified (e.g. B->C changes to B->C*), this change is not immediately transferred to the shortcut, leaving the old shortcut (A->C) to compete with the updated chain of associations (A->B, B->C*)

Together our findings demonstrate the effectiveness of using awake contextual TMR as a tool for examining the role of offline memory consolidation in shaping behaviour. We show that memory consolidation enhances inferential choice through the formation of shortcuts between indirect associations. However, these shortcuts cannot be rapidly updated. We thus demonstrate that memory consolidation during rest constructs a cognitive map that is more than the sum of its parts, allowing for efficient behaviour that is limited in flexibility.

## Results

### Task design and learning performance

To investigate the relationship between periods of rest and inference using TMR, we designed an inference task (Figure 1A-B). The inference task was adapted from a previous inference protocol implemented in humans and mice^22^ that leveraged a sensory preconditioning paradigm^2^. The inference task included three stages.

In the first stage, participants learnt to associate pairs of auditory and visual cues (‘associative learning’), with a many-to-one mapping from auditory to visual cues. The presentation of task cues was blocked into two groups, distinguished by different contextual background music that played throughout learning (either café or jungle music). One of these contextual background tracks was later played during rest to bias memory consolidation to this group of cues (TMR manipulation). After training, participants’ mean accuracy on the associative learning trials was 96.67% (SD=7.41%), with no significant difference in the maximum accuracy reached between sensory cues in the two contexts (café vs jungle; p=0.385) or between the group of cues that were subsequently subject to TMR and the group of cues that were not (TMR group vs no-TMR group; p=0.185) (Figure 1E).

In the second stage of the inference task, participants learnt to associate the visual cues with either a rewarding outcome (represented by a pound coin) or a neutral outcome (represented by a wooden coin) (‘conditioning’). As in the associative learning phase, contextual background music was played throughout learning. After training, participants achieved a mean accuracy of 99.50% on the conditioning trials (SD=2.50%) (Figure 1F).

Importantly, auditory cues were never paired with outcome cues, providing an opportunity to assess evidence for an inferred relationship between these indirectly related cues. Accordingly, in the final stage participants performed an ‘inference test’. In the inference test, we presented auditory cues in isolation, without visual cues or outcome cues, and without the contextual background music. In response to these isolated auditory cues, we measured evidence for inference by asking participants which outcome was associated with each auditory cue, despite the fact that they had never directly experienced this pairing.

### TMR manipulation

In addition to the inference task, we included a TMR manipulation to study the consequences of biasing memory consolidation. Participants underwent the TMR manipulation during a period of rest (60 minutes), after completing the associative learning and conditioning phases of the task, but before performing the inference test. Throughout this rest period, we played the contextual background track associated with cues in one of the two groups (i.e., either café music or jungle music, fully counterbalanced across participants) whilst participants completed a relaxing task unrelated to the learning task (namely, they completed a jigsaw puzzle). If TMR facilitates the construction of a cognitive map that is more than the sum of its parts, then this TMR manipulation should facilitate inference between cues in the TMR group.

### TMR improves inference performance

After the period of rest, participants performed the inference test (Figure 2A-B). Here, we presented the auditory cues in isolation and tested participants’ ability to infer a relationship with the corresponding outcome cue (rewarding or neutral). To assess the effect of TMR on inference, we compared participants’ accuracy in response to auditory cues in the TMR and no TMR groups.

**Figure 1.**
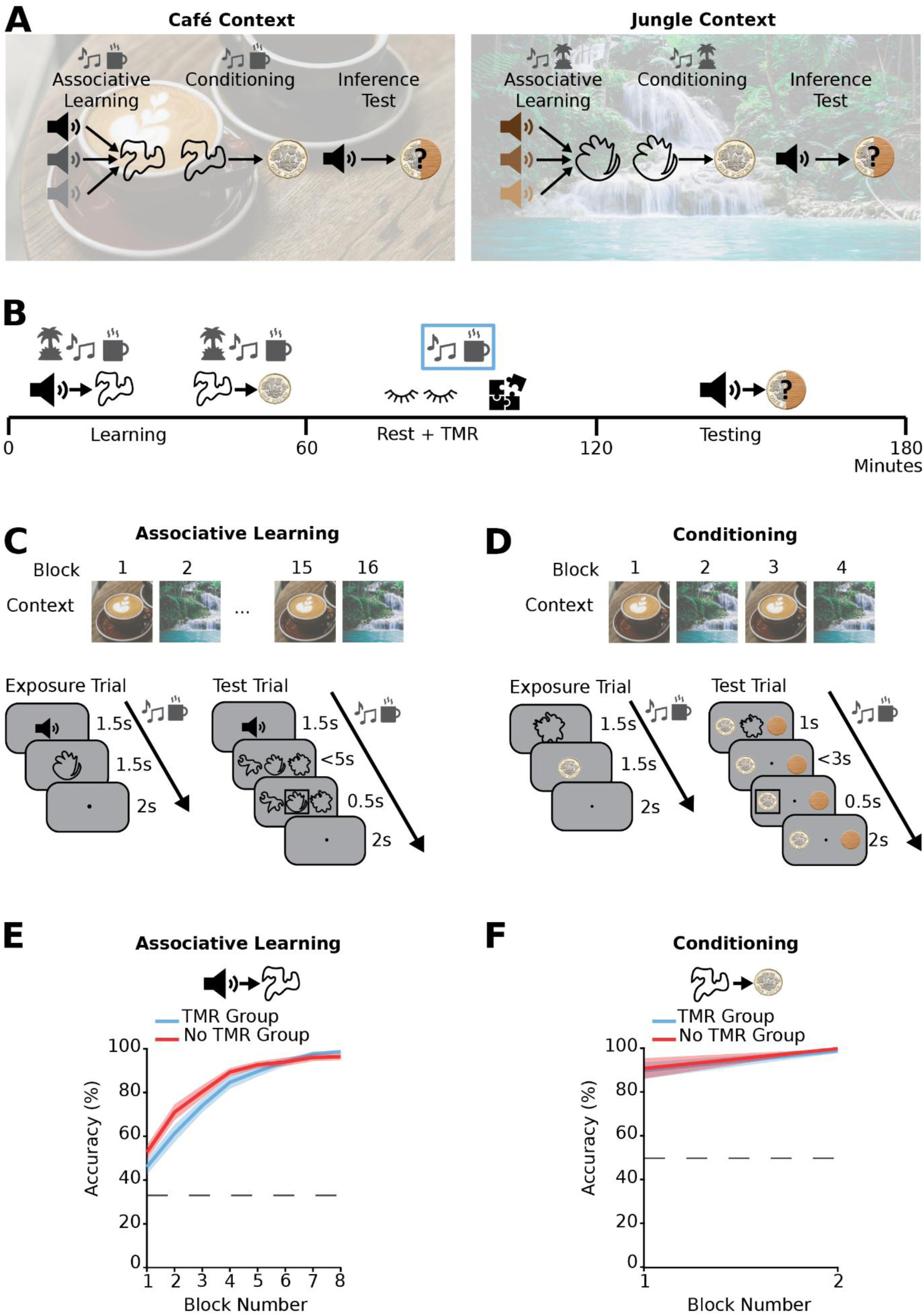
Inference Task Design and Learning Performance. (A) Three-stage inference task adapted from a previous protocol^22^. In the learning phase, participants learnt to associate auditory cues with visual cues (‘Associative Learning’), with a many-to-one mapping. Participants then learnt to associate visual cues with an outcome (rewarded – pound coin; neutral – wood coin) (‘Conditioning’). During the ‘Inference Test’, the auditory cues were presented in isolation, and we assessed participants’ ability to infer the value of any given auditory cue. Cues were divided into two sets, each grouped according to the contextual background music that played throughout learning (set 1: café context – left; set 2: jungle context – right). (B) Timeline of the task design in minutes. The task was split into three phases – learning, rest, and testing. During each block of the learning phase, the contextual background music for the relevant context was played. During the rest period, the contextual background music for only one set of cues was played to bias consolidation towards cues in this set (TMR group). After the rest period, participants were tested on their ability to infer associations between the auditory cues and the outcome cues. (C-D) Structure of the associative learning phase (C) and conditioning phase (D). Top: learning trials were blocked by context and alternated between the two contexts using the contextual background music. Bottom: Example trials (schematic). Blocks of trials consisted of both exposure trials (left) and test trials (right). (E-F) Behavioural performance during the associative learning (E) and conditioning (F) phases (mean accuracy ± SEM). Participants only proceeded to the inference test if they reached criterion (>85% performance accuracy for each group in both associative learning and conditioning). There was no significant difference in performance between cues in the two groups.

Participants were better at inferring the correct outcome for auditory cues in the TMR group (p=0.011) (Figure 2C). The inference task was carried out both in isolation, and in the presence of two distractor tasks (a one-back task and a semantic judgement task, see Methods). The distractor tasks were designed to increase the load on working memory or general cognitive resources respectively, to modulate task difficulty and avoid ceiling effects when inference was performed in isolation. In fact, participants were better at inferring the correct outcome for auditory cues in the TMR group under all three conditions (p=0.015, no distractor task; p=0.031, memory task; p= 0.030, non-memory task), with no significant difference in the TMR effect between conditions (p=0.130, no distractor task vs memory task; p=0.372, no distractor task vs non-memory task) (Fig. S1). Hence, hereafter results are pooled across the three distractor conditions. Overall, participants were better at inferring the correct outcome for auditory cues in the TMR group (p=0.011) (Figure 2C). This result demonstrates that biasing memory consolidation during periods of rest with awake, contextual TMR facilitates inferential choice.

### TMR creates a shortcut in memory between indirectly linked cues

Next, we asked *how* TMR influences the organisation of learned information to facilitate inferential choice. Specifically, we asked: do participants build a shortcut (A->C) that is separate from the original chain of learned associations (A->B, B->C), or do they instead strengthen the components of the learned associations (A->B, B->C)? We reasoned that if TMR facilitates inference by building a new short-cut between the auditory and outcome cues (A->C), the resulting cognitive map may be less amenable to flexible updating when associations between cues change.

Specifically, if the visual-outcome association B->C changes to B->C*, the correct chain of associations to support inference is now A->B and B->C*. If participants perform inference using the chain of learned associations (A->B, B->C), and each of the learned associations are merely strengthened during TMR, then it should be possible to rapidly update behaviour to account for the changes from B->C to B->C*. However, if an old shortcut (A->C) is represented that is not rapidly updated (to A->C*), behaviour may not account for the changes from B->C to B->C*.

To test this hypothesis, after the first inference test, we manipulated the association B->C (visual-outcome), such that the outcome associated with a subset of visual cues was reversed (’value flip’ manipulation, B->C*). Participants then performed the inference test once more, to test their ability to flexibly update the inferred value of auditory cues (from A->C to A->C*) (Figure 3A). Critically, if TMR creates a shortcut (A->C) between auditory and outcome cues that is separate to the chain of learned auditory-visual-outcome associations (A->B, B->C), then an update in the visual-outcome leg (B->C*) should not transfer to the auditory-outcome (A->C) short-cut. Therefore, to make correct inferences, changes to the visual-outcome (B->C*) mapping would require participants to draw on the complete chain of learned associations (A->B, B->C*) for cues in both the TMR and no TMR groups, negating the benefit of TMR. In addition, representation of an auditory-outcome (A->C) shortcut should compete with flexible updating in response to changes to the visual-outcome (B->C*) mapping.

**Figure 2.**
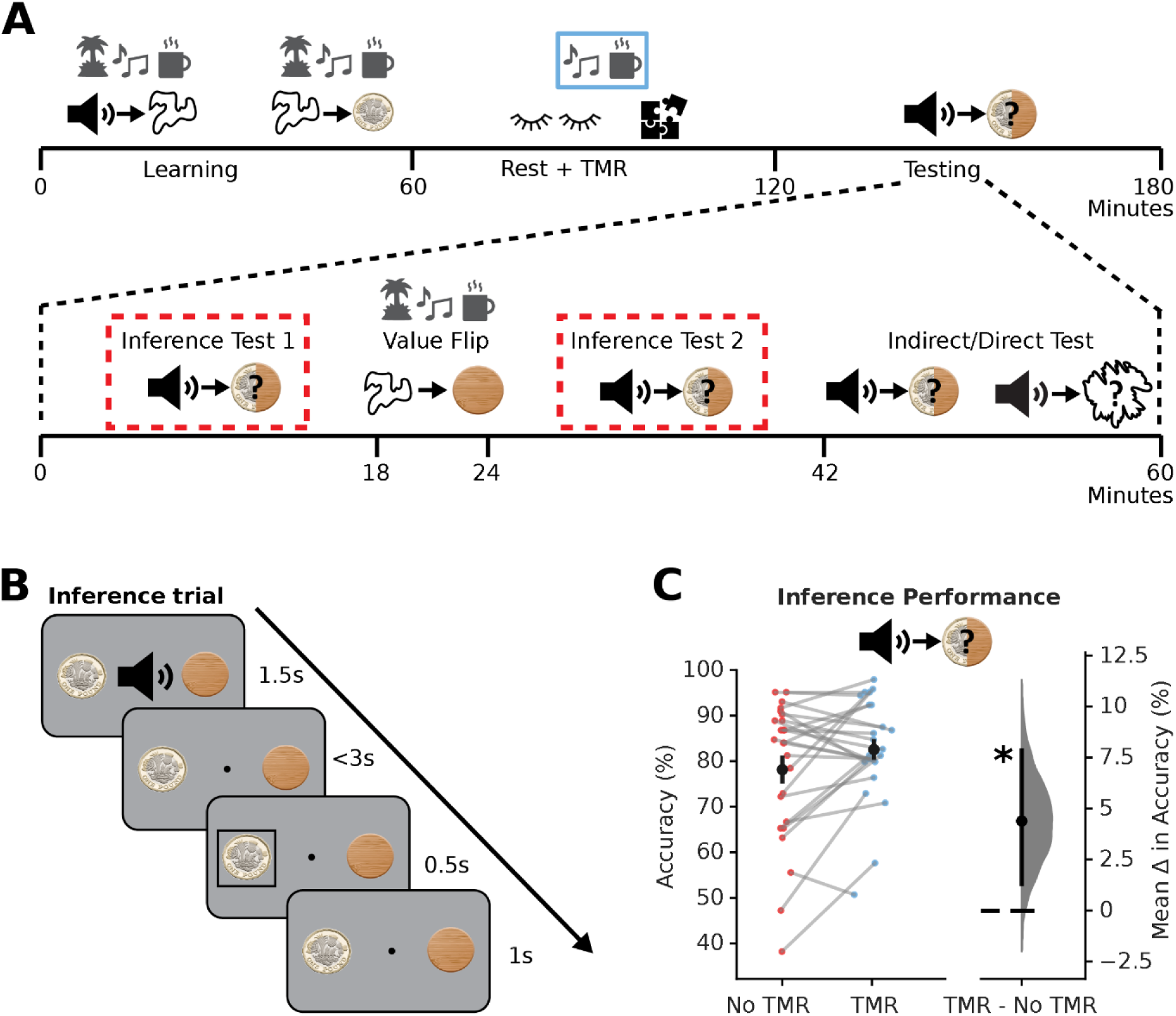
TMR Improves Inferential Choice. (A) Structure and timings of the testing phase of the task. Tests highlighted in red were combined to examine the effect of TMR on the ability to infer an association between the auditory cues and the outcome. The remaining stages (‘Value Flip’ and ‘Indirect/Direct Test’) are detailed in Figures 3 and 4. (B) Schematic: Example trial in the inference tests (I.e., ‘Inference Test 1’ and ‘Inference Test 2’). (C) Inference test performance. Left: raw data points for accuracy on auditory cues in the No TMR group (red; left) and TMR group (blue; right); each data point: mean accuracy for a given participant; black dot, mean; black ticks ± SEM. Right: difference in mean inference accuracy between auditory cues in the No TMR and TMR groups shown using bootstrap-coupled estimation (DABEST) plots^48^. Effect size for the difference between No TMR and TMR groups was computed from 10,000 bias-corrected bootstrapped resamples^52^: black dot, mean; black ticks, 90% confidence interval; filled-curve, sampling-error distribution. Participants were significantly better at inferring the correct outcome for auditory cues in the TMR group (p=0.011).

We observed evidence for both of these predicted effects. First, for visual-outcome mappings (B->C) where there was no change in the association (‘non-flipped’ associations), participants were better at inferring the correct outcomes for auditory cues in the TMR compared to the no TMR group (p=0.001). However, for flipped associations (B->C*), there was no significant difference between stimuli in the TMR and no-TMR groups (p=0.276) (Figure 3B-C). These results suggest that the benefit of TMR is negated by the value-flip manipulation. This was indeed the case, as the effect of TMR was significantly greater for non-flipped stimuli (p=0.024) (Figure 3D). This is consistent with the proposal that, after rapid updating in the visual-outcome (B->C*) mapping, participants had to perform inferential choice by drawing on the complete chain of learned associations (A->B, B->C*), rather than the shortcut (A->C), thus negating the benefit of TMR.

Next, we asked whether representing a shortcut between the auditory-outcome (A->C) cues acts in competition with the modified chain of learned associations (A->B, B->C*). Specifically, during the inference test performed after the value flip manipulation, we asked whether forming a shortcut (A->C) biases participants to look at the previously associated (but now incorrect) outcome (C) in the inference test. Thus, we used eye-tracking data acquired during the second inference test to ask whether, after the value flip manipulation, the shortcut (A->C) competes with the modified chain of associations (A->B, B->C*) (Figure 3E). To do this, we calculated the percentage of trials where participants looked at the incorrect outcome. For ‘flipped’ cues, the incorrect outcome corresponded to the outcome cue (C) that was previously correct prior to the value flip manipulation.

We hypothesised that for auditory cues where the visual-outcome association was flipped (B->C*), participants may still retain a shortcut (A->C) from the auditory cue to the (now incorrect) outcome for cues in the TMR group (Figure 3E) and would therefore be more likely to look at C during the decision process. This effect should be lower, or absent, for cues in the no-TMR group, as for these cues the shortcut (A->C) was weaker or not formed, relative to the TMR cues. This was indeed the case. For flipped cues, participants were significantly more likely to look at the incorrect outcome cue (C) in the TMR group compared to the no TMR group (p<0.001) (Figure 3D). This suggests that when a shortcut (A->C) forms due to TMR, knowledge of this shortcut competes with knowledge of relevant updated associations. This competitive effect supports the view that the shortcut (A->C) is separate from the chain of learned associations (A->B, B->C). Consequently, TMR may be considered to support efficient behaviour by facilitating inference through the formation of a shortcut between auditory and outcome cues even when these cues had not been directly experienced together. However, building a shortcut in the underlying cognitive map appears to have limitations – the shortcut cannot be flexibly updated in response to environmental change.

In a final set of analyses, we again used eye-tracking data to ask whether TMR facilitates inference by creating a shortcut in memory between the auditory and outcome cues. If a shortcut (A->C) is represented, we would expect participants to be less likely to use the intermediary visual cues (B) to help make inferential choices for cues in the TMR group. To test this, we analysed eye-tracking data acquired during a third inference test (‘indirect’) and during a memory test for learned associations (‘direct’) (‘indirect/direct’ test, Figure 2A, 4A-B). During the first half of this testing phase, participants again had to decide whether the auditory cues were associated with the rewarding or neutral outcome (inference test, ‘indirect’). Replicating our previous findings, we found that participants were again significantly more accurate at inferring the correct relationships for auditory cues in the TMR group compared to the no TMR group (p=0.013) (Figure 4E). However, unlike in the previous inference tests, all of the intermediary visual cues were now presented on the screen (Figure 4B). We predicted that if participants had formed a shortcut between auditory and outcome cues, they would be less likely to look at the visual cues to help them infer a value for each auditory cue. On correct trials, participants spent less time looking at the visual cues for auditory cues in the TMR group compared to the no TMR group (p=0.012) (Figure 4C). Therefore, for cues in the TMR group, there was a reduced need to look at the intermediary visual cue to help make a correct inference. Consistent with our previous findings, this result suggests that TMR facilitates the formation of a memory shortcut between the indirectly linked auditory and outcome cues.

**Figure 3.**
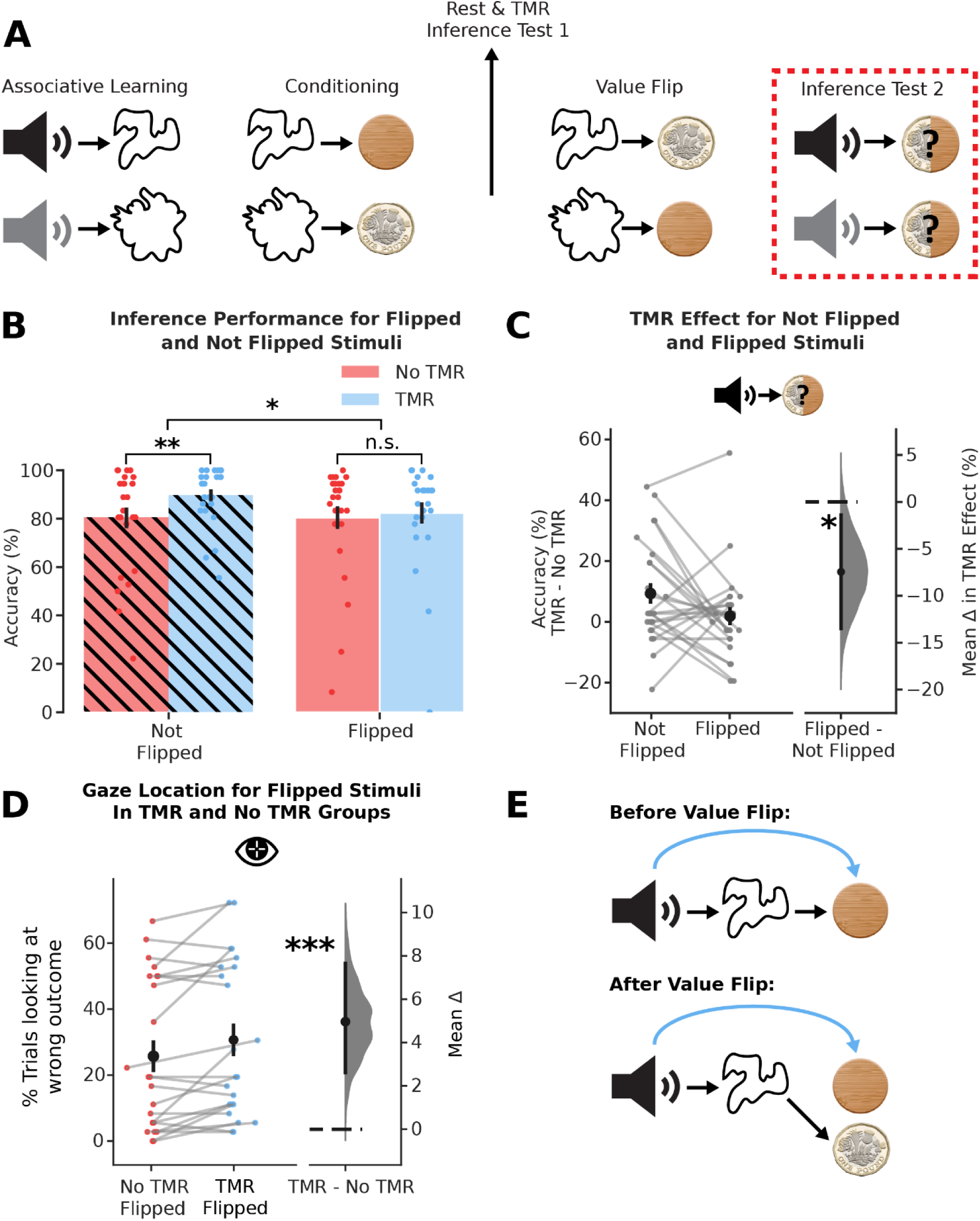
TMR Creates a Shortcut Which Cannot Be Readily Updated. (A) Schematic illustrating the value flipping manipulation. After the learning and rest phases, the outcome associated with half of the visual cues was flipped (‘Value Flip’). During Inference Test 2 (highlighted in red), we tested to see if participants could correctly infer the new outcome that was indirectly associated with the auditory cue. (B) Inference test performance after the value flip. Bars: mean percentage accuracy for No TMR (red; left two bars) and TMR (blue; right two bars) groups split by whether the visual cue-outcome association was not flipped (plain; left bar of each group) or flipped (hatched; right bar of each group); black ticks: ± SEM; each data point: mean accuracy for one participant. (C) Difference in TMR effect on inference test accuracy between auditory cues associated with not flipped and flipped. Left: raw data points for Not Flipped cues (left) and Flipped cues (right); each data point: mean effect of TMR (TMR accuracy – No TMR accuracy) for a given participant; black dots, mean; black ticks ± SEM. Right: difference in means between Not Flipped cues and Flipped cues shown using bootstrap-coupled estimation (DABEST) plots as in Figure 2C: black dot, mean; black ticks, 90% confidence interval; filled-curve, sampling-error distribution. The effect of TMR was significantly greater for the Not Flipped cues(p=0.024). (D) Difference in percentage of trials looking at the wrong outcome for stimuli where the visual cue-outcome association was flipped for the TMR and No TMR groups. Left: raw data points for No TMR group (red; left) and TMR group (blue; right); each data point: mean percentage of trials looking at wrong outcome for a given participant; black dots, mean; black ticks ± SEM. Right: difference in means between No TMR and TMR groups shown using bootstrap-coupled estimation (DABEST) plots as in Figure 2C: black dot, mean; black ticks, 90% confidence interval; filled-curve, sampling-error distribution. Participants spent a significantly higher percentage of trials looking at the wrong outcome for the TMR group (p<0.001). (E) Schematic showing how a shortcut may conflict with a chain of direct links after the value flip manipulation. Top: before value flip, both the chain of direct links (black arrows) and the shortcut formed by TMR (blue curved arrow) lead to the same outcome. Bottom: After value flip, the chain of direct links has been updated but now conflicts with the shortcut which cannot be rapidly updated.

### The effect of TMR on inference is not explained by improved memory for directly learnt associations

Finally, we asked whether awake contextual TMR also facilitates memory accuracy for directly associated cues. Previous work has shown that TMR protocols that use punctate auditory cues improve memory for directly learnt associations^31^. However, since our TMR protocol uses contextual cues, we wanted to investigate whether this protocol would have similar or different effects on memory. During the indirect/direct test (Figure 4A-B), participants were also given a memory test for learned associations between auditory and visual cues (‘direct associations’). By contrast to performance on the inferred relationships between auditory and outcome cues (‘indirect associations’) reported above (Figure 4E), when participants were tested on direct associations, they showed no significant difference in accuracy for auditory-visual (A->B) cue mappings in the TMR group compared to the no TMR group (p=0.478) (Figure 4F).

Since participants were trained to ceiling on the direct associations, it is possible that we did not see an effect of TMR due to ceiling performance. To investigate this potential confound, we performed a median split based on combined performance for TMR and no TMR groups, and applied this to both the indirect association (inference) test (in which we saw an effect of TMR) and direct association test (in which we did not see an effect of TMR). For direct associations, even for participants who were not at ceiling, we continued to observe no significant effect of TMR (p=0.451, top 50%; p=0.442, bottom 50%), suggesting that the lack of improvement under TMR was probably not due to a ceiling effect. In contrast, for indirect associations (where we had observed an overall effect of TMR), the TMR effect remained significant for both the bottom half (p=0.029) and top half (p=0.036) of participants. Thus, even for participants who performed the inference task close to ceiling performance, we continued to observe a significant effect of TMR. Together these findings suggest that the reported difference in effect of TMR on indirect and directly learned associations is likely not due to difference in task difficulty or due to a ceiling effect, but instead indicates that contextual TMR applied during rest prioritises formation of shortcuts that support inference, rather than strengthening learned memory content.

Together, these results suggest that while contextual TMR improves inferential choice across indirectly associated cues, this improvement cannot be explained by improved performance on directly learned associations. Therefore, the benefits of contextual TMR on inferential choice must be explained by a mechanism that goes beyond merely strengthening learned associations. This provides further evidence to suggest that periods of rest facilitate inference by restructuring knowledge and creating shortcuts between items in memory that extend beyond direct experience.

**Figure 4.**
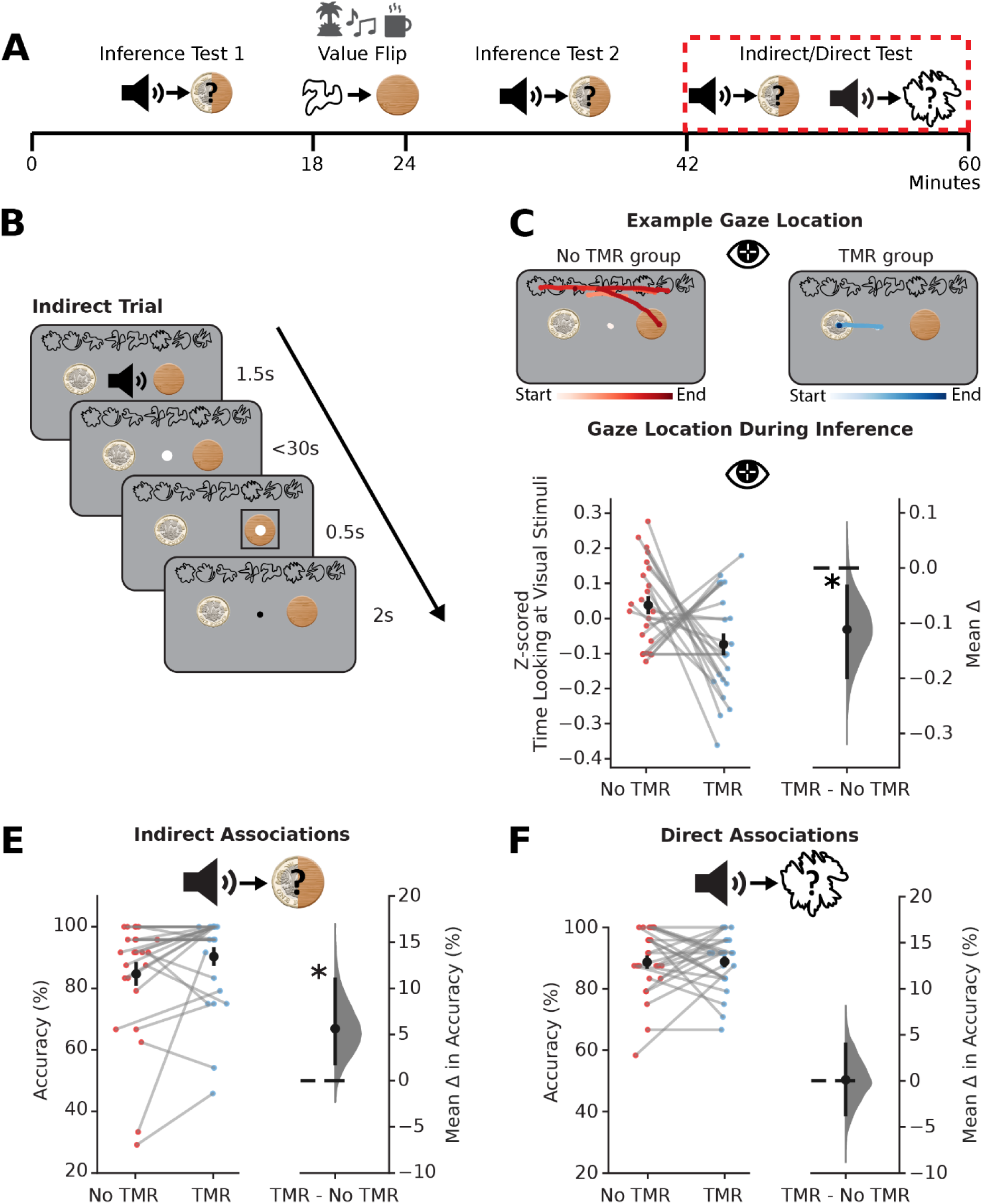
TMR Creates a Shortcut Which Does Not Rely on Directly Learnt Associations. (A) Structure of the testing phase of the task. Data presented in (C-F) is from the ‘Indirect/Direct Test’ highlighted in red. In the indirect/direct test, participants were tested on both the indirect (auditory – outcome) and direct (auditory – visual) associations for each auditory cue. (B) Schematic: example indirect trial (schematic) where auditory cues were now presented while the visual cues were on screen. Participants were required to infer the outcome cue in response to the auditory cue. Direct trials followed the same structure except, after hearing the auditory cue, participants had to choose the associated visual cue rather than the associated outcome. (C) Gaze location from eye tracking data acquired during indirect trials in the indirect/direct test. Top: example gaze trajectories from a single indirect trial for one participant in response to an auditory cue from the No TMR group (red; left) and an auditory cue from the TMR group (blue; right). Darker colours indicate later time in the trial. Bottom left: raw data points for No TMR group (red; left) and TMR group (blue; right); each data point: mean z-scored time spent looking at visual cues during inference (correct trials only); black dots, mean; black ticks ± SEM. Bottom right: difference in means between No TMR and TMR groups shown using bootstrap-coupled estimation (DABEST) plots as in Figure 2C: black dots, mean; black ticks, 90% confidence interval; filled-curve, sampling-error distribution. Participants spent significantly less time looking at the visual cues in response to auditory cues in the TMR group (p=0.012). (D-E) The effect of TMR on indirect associations (D) and direct associations (E). Left of each panel: raw data points for No TMR group (red; left) and TMR group (blue; right); each data point: mean accuracy for a given participant; black dots, mean; black ticks ± SEM. Right of each panel: difference in means between No TMR and TMR groups shown using bootstrap-coupled estimation (DABEST) plots as in Figure 2C: black dot, mean; black ticks, 90% confidence interval; filled-curve, sampling-error distribution. There was a significant effect of TMR on accuracy for indirect associations (p=0.013) but not for direct associations (p=0.478).

## Discussion

Previous studies have shown the benefit of rest/sleep for inference^5–9,32,33^. However, the nature of the changes that occur during rest/sleep to facilitate inference remain unclear. In this study, the central question we ask is: *<u>How</u> does memory reactivation during rest reorganise memory to guide adaptive behaviour?* We sought to tease apart two ways in which rest/sleep could affect memories: an improvement in behavioural performance on an inference task could be supported by either a quantitative strengthening of learned, component associations (A->B and B->C), or a qualitative change, involving the formation of a new, ‘shortcut’ link (A->C). The latter, qualitative change would reflect an essential pre-requisite for the organization of knowledge into a structured cognitive map that is more than the sum of its parts.

Using a multi-stage inference task with awake auditory contextual TMR, we show that memory reactivation during rest benefits inference by building a new shortcut (e.g. A->C), with no effect on directly trained associations (e.g. A->B, B->C). Thus, promoting memory consolidation during rest facilitates the formation of novel shortcuts in the cognitive map. This may explain why periods of rest and sleep facilitate behavioural readouts of inference, but also other cognitive processes such as insight, generalisation and abstraction^5–10^. Interestingly, both our behavioural and eye-tracking data show that these shortcuts cannot be readily updated in response to subsequent changes in the environment. Rather, shortcuts (e.g. A->C) may limit behavioural flexibility when optimal behaviour requires participants to revert to a chain of learned associations (i.e. A->B, B->C). Together, our results reveal how memory reactivation during rest plays a causal role in *restructuring* memories, while highlighting how the organisation of knowledge sets an important trade-off between efficient and flexible behaviour.

A candidate neural mechanism that may explain our results is replay. During an inferential learning task in mice, similar to the one employed here, replay was shown to go beyond direct experience by “joining-the-dots” between indirectly linked cues^22^. During awake sharp-wave ripples (SWRs), hippocampal spiking activity represented the inferred relationship (A->C) in the absence of the intermediary cue (B). However, the behavioural consequences of this form of replay were not investigated. Here in humans, we use a TMR manipulation to bias the content of replay^14^. Unlike previous TMR protocols, we used awake, contextual TMR to demonstrate that biasing memory consolidation during rest leads participants to form a shortcut between indirectly related cues. Furthermore, this shortcut reduces the need to rely on memory for the intermediary cue. Together these findings suggest that our TMR manipulation facilitates replay of inferred relationships between cues in the TMR group, promoting the formation of a shortcut in the underlying cognitive map. However, since this study did not directly measure replay, future investigations will need to validate this interpretation using neuronal recordings.

While our ability to make novel inferences can clearly support efficient behaviour, representing inferred relationships as a shortcut (A->C) rather than a chain of learned associations (A->B, B->C) may limit behavioural flexibility. Our data demonstrates the limitation of building a short-cut in two key analyses. These two analyses are applied to data acquired after the value flip manipulation, where half of the visual-outcome mappings are ‘flipped’ (B->C becomes B->C*). First, we show that the TMR benefit on performing inference is no longer observed for the flipped cues. Second, for the flipped cues, eye-tracking data shows that participants spend more trials looking at the previously correct, but now incorrect outcome (C*) for TMR compared to no-TMR cues. This suggests that the outdated shortcut (A->C) may compete with the updated chain of associations (A->B, B->C*). As well as supporting the proposal that the shortcut (A->C) is indeed separate from the chain of associations (A->B->C/C*). These data demonstrate that shortcuts in memory (e.g. A->C) cannot be readily updated in response to rapid changes in the environment and may limit behavioural flexibility when optimal behaviour requires participants to revert to a chain of learned associations (i.e. A->B, B->C*). We speculate that if a further rest period was included after the change in value of outcome (C), a new updated shortcut (A->C*) would have been formed and used. Indeed, there is evidence to suggest that rest and sleep may play a role in integrating new memories into existing knowledge^34–37^. Our findings complement this work by suggesting that adaptive re-organisation of the cognitive map cannot always occur ‘on the fly’ and may instead rely on offline periods of rest or sleep. Our study therefore provides insight into how new information is incorporated into a cognitive map, to guide adaptive behaviour.

The above findings were made possible by our novel contextual auditory TMR protocol. The majority of previous TMR studies have used either punctate auditory cues^31^ or contextual odour cues^38^. Using punctate auditory cues may have several drawbacks, including the need to precisely time the cue to specific phases of sleep oscillations to see the best effects^39^. Furthermore, since punctate cues are associated with specific stimuli, they provide a limited tool to investigate how links across multiple stimuli are consolidated. The contextual TMR protocol presented here provides a unique opportunity to investigate the effect of memory reactivation across an entire map of associations. We demonstrate how memory reactivation during periods of rest influences the organisation of memories within the cognitive map, where memory reactivation plays a causal role in forming novel links that go beyond direct experience.

An additional benefit of the TMR protocol used here is that it is very simple to implement. Using a contextual background track removes the need to present multiple different cues across the rest period. Furthermore, auditory cues are simpler to deliver than the contextual odour cues that have been used in previous contextual TMR paradigms. Finally, delivering the TMR during awake rest, as opposed to sleep, removes a significant time investment and the need to ensure for accurate sleep staging. Given the increasing interest in using TMR for clinical benefit in a number of neuropsychiatric conditions^40^, the simplicity of the paradigm carries important translational potential.

Previous work, which largely focus on the effect of punctate TMR cues applied to directly experienced associations, show variable results when using TMR during periods of awake rest^26,31,38,41,42^, which may in part be explained by variation in task difficulty and performance accuracy. Indeed, non-invasive measures of memory reactivation suggest that replay of directly learned information prioritises weakly learned information^43^. Consistent with this view, TMR facilitates direct associations by reducing forgetting^31,44,45^. Therefore, by prioritising reactivation of weaker memories, TMR may help protect memories that are most vulnerable to forgetting^43^. In our data we report a specific benefit of TMR for indirect associations, with no effect found on direct associations. In our protocol we included only a short time interval (1 hour) between learning and test, which may have reduced the scope for forgetting. It is possible that if we had used a longer time interval, we would have seen a benefit of TMR for directly learned associations. Notwithstanding this point, the lack of an effect on direct associations implies that the observed restructuring of knowledge does not rely on strengthening of directly learned associations, or on rescuing the direct associations from forgetting.

We also note that the differential effect of TMR on indirect and direct associations cannot easily be explained by differences in performance accuracy or ceiling effects. Unsurprisingly, participants showed higher performance accuracy on direct compared to indirect associations. However, when we performed a median split analysis using the combined performance for TMR and no TMR cues, we show that both high and low performing participants continue to show a significant TMR effect for indirect but not direct associations. Therefore, participants who showed performance accuracy close to ceiling continued to show a significant TMR effect on indirect associations, while participants where performance was not at ceiling continued to show no significant effect of TMR on direct associations. Together these findings suggest that the differential effect of TMR on direct and indirect associations cannot be explained by differences in performance accuracy. Nevertheless, indirectly trained associations may share physiological properties with weaker memories, rendering them more susceptible to TMR.

Overall, our results demonstrate that knowledge is subject to qualitative restructuring during offline periods of rest/sleep. During rest, stimuli that are reactivated (due to contextual TMR) are organized into cognitive maps through the creation of novel shortcuts between indirectly linked cues. These shortcuts are distinct from the original chain of learned associations and the behavioural benefit of forming a shortcut cannot be explained by quantitatively strengthening the component learned associations (since there was no benefit of TMR on these learned associations even in participants whose performance was well below ceiling). Furthermore, when the component learned associations are rapidly modified in response to changes in the environment, shortcuts are not updated. Indeed, an outdated shortcut can conflict or compete with an updated chain of associations to control decision making, illustrating that the two sets of associations are separate. Together our findings provide new insight into how integrated cognitive maps are formed during periods of rest. Moreover, we demonstrate the effectiveness of using awake contextual TMR as a tool to reveal how the resting brain re-organises memory to shape adaptive behaviour.

## Methods

### Participants

A total of 32 healthy participants were recruited for this study. 7 participants were excluded from analysis as they failed to reach the performance criterion during the initial associative learning phase (criterion: >85% performance accuracy for each contextual group of cues). The remaining 25 participants were included in the analysis (mean age of 24.72 ± 6.35 years, range 19-40, 4 males). All participants had normal or corrected-to-normal vision. All experiments were approved by the University of Oxford ethics committee (reference number: R43593/RE013).

### Experimental set-up

Cues were presented on a 24-inch screen with a spatial resolution of 1920 x 1080 pixels (width x height), a background luminance of 0.5 (grey), and a refresh rate of 100 Hz. The approximate distance of participants from the screen was 64cm, which meant that one degree of visual angle corresponded to 40 pixels on the screen. Stimulus presentation was controlled using Psychophysics Toolbox-3^46^ in MATLAB (version R2015b). Eye movements were recorded in the testing phase of the experiment (after the rest period) using an eye-tracking camera (EyeLink®, SR Research) tracking both eyes at a rate of 1kHz.

### Inference task - overview

Participants performed an inference task (Figure 1A-B). In the first stage of the task (‘associative learning’), participants learnt to associate punctate auditory cues with visual cues. The punctate auditory cues were naturalistic sounds from the BBC sound library (https://sound-effects.bbcrewind.co.uk/) and the visual cues were unsymbolic shapes^47^. Each visual cue was paired with three auditory cues. In the second stage of the task (‘conditioning’), these visual cues were then associated with either a pound coin (rewarded outcome) or a wood coin (neutral outcome, of no value). The cues were split into two sets, grouped according to the contextual background music that played throughout learning. For one of the two sets of cues, participants heard jungle music in the background and the punctate auditory cues were sounds you would hear in a jungle (e.g., animal noises). For the other group, participants heard café music in the background and the punctate auditory cues were sounds you would hear in a café (e.g., a cash register). In total, 12 auditory cues and 4 visual cues were included in each group and the relationship between the conditions (i.e., which context was learnt first, and which context was subject to TMR) was fully counterbalanced across participants. Since the visual cues had no direct meaning, they were randomly assigned to each group across participants.

### Behavioural protocol

The associative learning stage was designed to allow participants to learn associations between auditory and visual cues. This stage of the task included passive exposure and testing trials (Figure 1C). In the exposure phase, on each trial a punctate auditory cue was presented for 1.5s, immediately followed by presentation of a visual cue for 1.5s, followed by an inter-trial interval (ITI) of 2s. Each mini-block of exposure consisted of 12 trials (one presentation of each auditory-visual pair). Each mini-block of exposure was followed by a testing mini-block. On each trial in the testing mini-block participants heard an auditory cue for 1.5s before being presented with 3 possible visual cues. Participants were then required to select the visual cue that was paired with the auditory cue they just heard, responding within 5s using the keys ‘j’, ‘k’, and ‘l’. Once a choice had been selected, a black rectangle would highlight this choice for 0.5s, followed by an ITI of 2s. At the end of each testing mini-block, participants were given feedback on their average performance. Each block in the associative learning phase was made up of one exposure mini-block followed by one testing mini-block and consisted only of cues from one of the two groups of cues, with the relevant contextual background music playing throughout the block. Blocks alternated between the two contexts with a total of 8 blocks per context. The relationship between the conditions (i.e., which context was learnt first, and which context was subject to TMR) was fully counterbalanced across participants.

The conditioning stage was designed to allow participants to learn associations between visual and outcome cues. The blocks followed the same structure as the associative learning stage, including both passive exposure and testing trials (Figure 1D). In each trial of the exposure phase, a visual cue was first presented for 1.5s, immediately followed by the associated outcome for 1.5s, followed by an ITI of 2s. Each exposure mini-block consisted of 4 trials (one presentation of each visual cue), which was followed by a testing mini-block. In each trial of the testing mini-block, a visual cue was presented together with the two outcome cues for 1s. After this 1s, the visual cue would disappear but the outcome cues would remain on the screen and the participants had a further 3s to make their choice (total of 4s response period). Participants had to select which outcome they thought was associated with the visual cue using the keys ‘j’ and ‘l’. Once a choice had been selected, a black rectangle would highlight their choice for 0.5s, followed by an ITI of 2s. Participants were given feedback on their average performance at the end of each mini-block of 4 trials. Each block in the conditioning phase was made up of one exposure mini-block followed by one testing mini-block and consisted only of cues from one of the two groups of cues, with the relevant contextual background music playing throughout the block. Blocks alternated between each context with a total of 2 blocks per context. The relationship between the conditions (i.e., which context was learnt first, and which context was subject to TMR) was fully counterbalanced across participants.

The TMR manipulation was performed after the conditioning stage. During TMR, participants were instructed to rest for up to 60 minutes, during which participants were asked to alternate between resting with their eyes closed for 10 minutes and doing a jigsaw puzzle for 20 minutes. In total, they spent 20mins with their eyes closed and 40mins doing a jigsaw puzzle. During this rest period, the contextual soundtrack associated with the TMR group was played throughout. The contextual soundtrack used for TMR was fully counterbalanced across participants. To encourage participants to attend to the soundtrack, the soundtrack was turned on and off for periods of between 9 and 16s, alternating with periods of silence. The length of each on and off period was selected from a random distribution.

After the TMR manipulation, participants were required to perform an inference test. During the inference test, participants were asked to infer the relevant outcome in response to each auditory cue.

Before beginning the first inference test, participants practised performing two tasks together using only two auditory cues, not taken from the task, and a 1-back task using digits rather than shapes. The practice consisted of up to 4 blocks with 24 trials per block. The timings of these practice trials matched the timings of the main inference test described below.

We designed three different versions of the inference test which included distractor tasks designed to increase the load on visual working memory or general cognitive resources (Figure S1A). For all versions, participants were presented with an auditory cue and had to choose which of the two outcomes they thought was associated with the auditory cue (Figure 2B). The auditory cue was presented together with the two outcome cues for 1.5s. After 1.5s, the auditory cue would stop but the outcome cues would remain on the screen and the participants had a further 3s to make their choice (total of 4.5s response period). Participants had to select which outcome they thought was associated with the auditory cue using the keys ‘j’ and ‘l’. Once a choice had been selected, a black rectangle would highlight this choice for 0.5s, followed by an ITI of 1s. Participants were told they would receive a bonus reward of up to £5 based on how many cues they correctly inferred. Participants were given feedback on their average performance at the end of each block. Auditory cues from both contextual groups were randomly interleaved in each block and there was no background music during the inference tests.

For 2 of the 6 blocks, the participants also performed a visual memory distractor task (Figure S1A). The purpose of this distractor task was to modulate task difficulty: a visual memory distractor task could impair recall of the intermediate visual cue when participants perform inference. The visual memory distractor task involved using a 1-back task with shapes similar to those seen in the task. On each trial, between presentations of each auditory cue, a shape would appear for 2s, and the participants had to respond based on whether it was the same or different from the previous shape using the keys ‘a’ and ‘d’. The shape would become darker (participant chose ‘same’) or lighter (participant chose ‘different’) after the participant had made their choice. The shape remained on screen until the 2s was complete. The shapes would appear in a random orientation which participants were instructed to ignore. Following a pause of 1s, the auditory cue would begin, and the rest of the inference trial would commence as described above.

For another 2 of the 6 blocks, participants performed a non-memory-based distractor task (Figure S1A). The purpose of this distractor task was again to modulate task difficulty, but without the interfering effect of a visual memory load. On each trial, between presentations of each auditory cue, an image would appear for 2s, and the participants had to respond based on whether the image depicted something living (e.g., a hedgehog) or something non-living (e.g., balloons). The image would become darker (participant chose ‘living’) or lighter (participant chose ‘non-living’) after the participant had made their choice. The image remained on screen until the 2s was complete. Following a pause of 1s, the auditory cue would begin, and the rest of the inference trial would commence as described above.

For the remaining 2 blocks, there was no distractor task. After the normal ITI of 1s, there would be an additional pause of 3s to ensure trials across all versions of the inference task were the same length. Following this pause, the auditory cue would begin, and the rest of the inference trial commenced as described above. These 2 blocks which did not include distractor tasks always occurred at the end to minimise order effects of participants practising the inference task on its own.

After all 6 blocks of the first inference test, participants redid the conditioning stage of the experiment but with half of the associations between visual and outcome cues flipped (‘value flip’) (Figure 3A). The protocol for the value flip stage was otherwise the same as for the conditioning stage (Figure 1D) except participants completed 1 block per context (rather than the 2 blocks per context done in the initial conditioning stage). After the value flip, a second inference test (‘inference test 2’) was conducted using the same procedure as previously described and also included 6 blocks (Figure 2B; Figure S1A).

In the indirect/direct test, we tested both the indirect (auditory–outcome) and direct (auditory-visual) associations paired with each auditory cue (Figure 4B). All visual cues and outcomes were present on the screen. By having all cues on the screen, we could test whether there were differences in how long participants spent looking at the visual cues during inference in the TMR and no TMR groups. For the first 2 blocks, the auditory cue was presented, and participants had to select one of the two outcomes using the mouse. For the final 2 blocks, the auditory cue was presented, and participants had to select one of the 8 possible visual cues which was paired with the sound. This allowed us to assess performance on the directly learnt associations as well as the indirect auditory-outcome association. For all blocks, the auditory cue was presented for 1.5s. Following this, participants had 30s to select either the outcome cue (first 2 blocks) or the visual cue (last 2 blocks) associated with the auditory cue. Participants made their selection by moving the cursor over the chosen cue and pressing ‘spacebar’ to confirm. Once a choice had been selected, a black rectangle would highlight this choice for 0.5s, followed by an ITI of 1s. Participants were given feedback on their performance at the end of each block.

### Eye-tracking Data Preprocessing

For 2 participants, no usable eye-tracking data was collected. The remaining 23 participants were all included in eye-tracking analyses. The conversion of the EyeLink® 1000 Edf files was done with the Edf2Mat Matlab Toolbox designed and developed by Adrian Etter and Marc Biedermann at the University of Zurich. Gaze position and pupillometry data were smoothed using a 50ms Gaussian kernel. Blinks were detected from the smoothed pupillometry data and removed from the gaze position data. For each participant, the eye with the fewest missing samples was selected and used for all further analyses. The gaze position was epoched around the auditory cue onset for trials in the second inference test and for indirect trials in the indirect/direct test. For the comparison of flipped cues, we analysed the percentage of trials where participants looked at the incorrect outcome cue. To investigate whether participants relied on knowledge of the intermediary visual cue, we analysed the time spent looking at the visual cues in response to auditory cues in the TMR and no TMR groups on correct trials of the indirect test. Due to the large variation in the average time spent looking at the visual cues, we z-scored the fixation lengths across all trials for each participant.

### Analysis

Analysis was conducted in MATLAB (version R2022a) and in Python v.3.6 (https://www.python.org/downloads/release/python-363/), using the Python packages DABEST^48^, scipy^49^, numpy^50^, matplotlib^51^, seaborn^52^, pandas^53^. Throughout this study, we used a bootstrap-coupled estimation of effect sizes^54^. For each estimation plot showing a difference between TMR and No TMR groups: the left panel shows the distribution of raw data points for the entire dataset, superimposed on plots reporting group means ± SEM; and the right panel displays the difference between the TMR group and the No TMR group, computed from 10,000 bootstrapped re-samples. For each estimation plot: black dot, mean; black ticks, 90% or 95% confidence interval; and filled curve: bootstrapped sampling error distribution. We used 90% confidence intervals (one-sided) for tests of difference between TMR and No TMR groups. For any other tests, we used 95% confidence intervals (two-sided).

## Data and Code Availability Statement

The data and code used in this study will be made available via the MRC BNDU Data Sharing Platform (https://data.mrc.ox.ac.uk) upon publication.

## Acknowledgements

This research was supported by funding from the Medical Research Council (UKRI-MRC) (MR/W01971X/1, MR/L019639/1 to J.X.R.), UKRI (MR/W008939/1 to H.C.B.), and the Biotechnology and Biological Sciences Research Council (UKRI-BBSRC) (BB/M011224/1 to A.B.R.). The Wellcome Centre for Integrative Neuroimaging is supported by core funding from the Wellcome Trust (203139/Z/16/Z). The funders had no role in study design, data collection and analysis, decision to publish or preparation of the manuscript.

## Author Information

### Authors and Affiliations

**Department of Experimental Psychology, University of Oxford, Oxford, United Kingdom**

Cal M. Shearer, Jill X. O’Reilly

**Medical Research Council Brain Network Dynamics Unit, Nuffield Department for Clinical Neurosciences, University of Oxford, Oxford, United Kingdom**

Cal M. Shearer, Annalise B. Rawson, Helen C. Barron

**Wellcome Centre of Integrative Neuroimaging, University of Oxford, FMRIB, John Radcliffe Hospital, Oxford, United Kingdom**

Helen C. Barron, Annalise B. Rawson

### Contributions

All authors contributed to the experimental design and editing the paper. C.M.S, H.C.B, and J.X.R contributed to data analysis and writing the paper. C.M.S. collected the data.

### Corresponding author

Cal Shearer

## Ethics declarations

### Competing interests

The authors declare no competing interests.

## Supplemental Information

**Figure S1.**
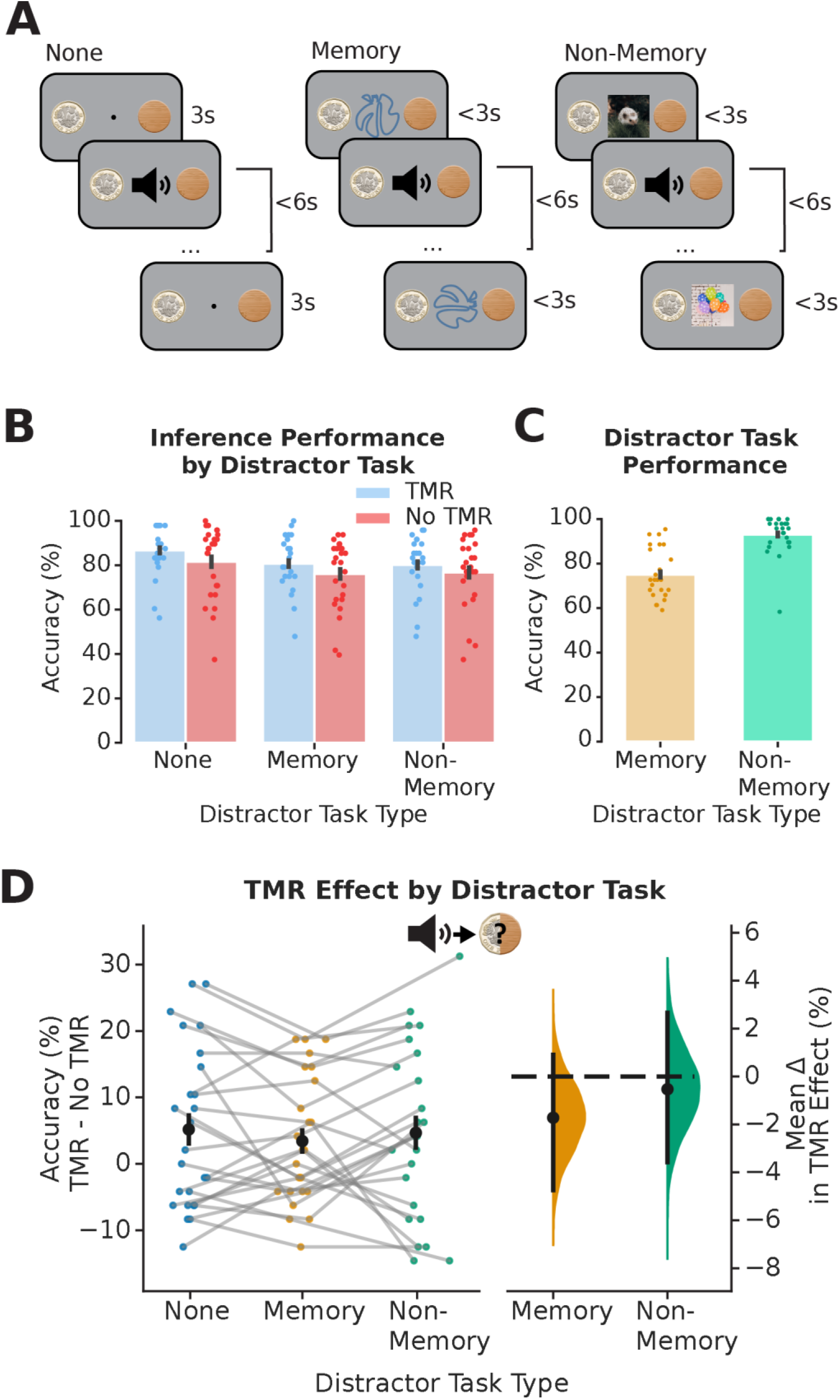
Distractor Task Does Not Change the Effect of TMR On Inferential Choice. (A) Schematic: Example trials for different distractor task types during the inference test. Trials start with either no distractor task (left), a memory distractor task (participants are asked if a shape is the same or different to the one seen on the previous trial (middle)), or a non-memory secondary task (participants are asked to identify if a photo is of something living or non-living (right)). On all trial types, participants then hear an auditory cue and are asked to infer the outcome associated with that auditory cue. Full details of the inference part of each trial are shown in Figure 2B. (B) Inference test performance broken down by distractor task type and TMR group. Bars: mean percentage inference accuracy for no distractor task (left group), memory task (middle group), and non-memory task (right group) groups split by TMR (blue; left bar of each group) and No TMR (red; right bar of each group); black ticks: ± SEM; each data point: mean accuracy for one participant. (C) Distractor task performance. Bars: mean percentage accuracy on the distractor task for memory task (orange; left) and non-memory task (green; right); black ticks: ± SEM; each data point: mean accuracy for one participant. (D) Difference in TMR effect on inference between different distractor tasks. Right: raw data points for no distractor task (blue; right), memory task (orange; middle), non-memory task (green, right); each data point: mean effect of TMR (TMR group accuracy – No TMR group accuracy) for a given participant; black dots, mean; black ticks ± SEM. Right: difference in means between no distractor task (dashed black line) and other distractor task conditions shown using bootstrap-coupled estimation (DABEST) plots as in Figure 2C: black dots, mean; black ticks, 95% confidence interval; filled-curve, sampling-error distribution. There was no significant difference in the effect of TMR on accuracy between distractor task types (p=0.130, no distractor task vs memory task; p=0.372, no distractor task vs non-memory task).

**Figure S2.**
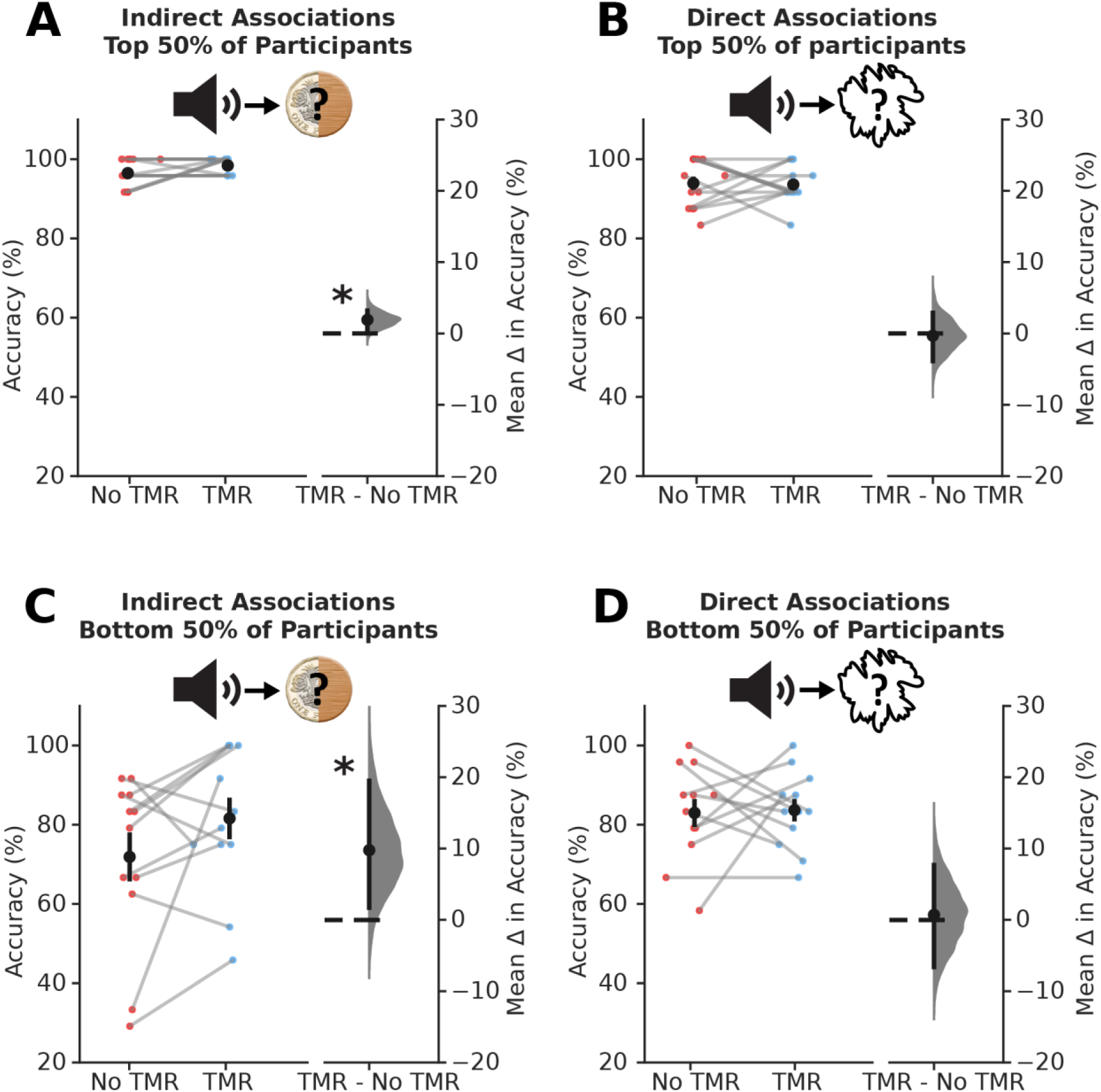
Differences in Overall Accuracy Cannot Explain the Effect of TMR on Direct and Indirect Associations. (A-D) The effect of TMR on indirect associations (A; C) and direct associations (B; D) for participants split by overall accuracy in the top 50% (A-B) and bottom 50% (C-D). Left of each panel: raw data points for No TMR group (red; left) and TMR group (blue; right); each data point: mean accuracy for a given participant; black dots, mean; black ticks ± SEM. Right of each panel: difference in means between No TMR and TMR groups shown using bootstrap-coupled estimation (DABEST) plots as in Figure 2C: black dot, mean; black ticks, 90% confidence interval; filled-curve, sampling-error distribution. There was a significant effect of TMR on accuracy for indirect associations (i.e. inference) for both the bottom (p=0.029) and top (p=0.036) 50% of participants. There was no significant effect of TMR on accuracy of direct associations, for either the bottom of top 50% of participants (p=0.451, top 50%; p=0.442, bottom 50%)

## Notes

### Competing Interest Statement

The authors have declared no competing interest.

### Summary of Updates

New analyses and revisions to the text made in response to comments made by a journal editor. Figure 3 revised; Figure S2 added.

